# Chromatin mimicry by human JC virus

**DOI:** 10.1101/2024.11.04.621823

**Authors:** Uwe Schaefer, Yekaterina A. Miroshnikova, Wei Xie, Adam G. Larson, Ziyu Lu, Shoudeng Chen, Martina Bradic, Yehuda Goldgur, Kexin Chen, Ved P. Sharma, Junyue Cao, Dinshaw J. Patel, Geeta J. Narlikar, Sara A. Wickström, Alexander Tarakhovsky

## Abstract

Chronically persistent viruses are integral components of the organismal ecosystem in humans and animals ^1^ ^2^. Many of these viruses replicate and accumulate within the cell nucleus ^3^. The nuclear location allows viruses to evade cytoplasmic host viral sensors and promotes viral replication ^4^. One of the unexplored and puzzling aspects of the viral nuclear lifecycle involves the virus’s ability to deal with the physical constraints of nuclear architecture. To replicate and accumulate within the nucleus in large numbers sufficient for infection spreading, DNA viruses need to overcome the spatial limitations imposed by chromatin and the nuclear matrix. We found that one of the most widespread and potentially lethal human viruses, the JC polyomavirus ^5^, interferes with nuclear heterochromatin to create virus-occupied space. The JC virus’s impact on heterochromatin is mediated by the viral nonstructural protein, Agnoprotein (Agno). Agno’s interference with heterochromatin is governed by structurally diverse mimics of host epigenetic regulators that facilitate virus-induced chromatin reorganization and a dramatic decline in nuclear stiffness in the infected cells. The JCV epigenetic mimicry is critical for the virus infection, as evident from reduced replication of mimic-mutant viruses. Our data suggest that modulation of nuclear mechanical properties is a novel strategy enabling chronicity of the JC and possibly other nuclear virus infections.

---

The JC virus (JCV), which replicates and accumulates in the nuclei of infected cells ^6^, infects approximately 70-80% of the human population in the Western world ^7 8 9^, typically in a chronic and largely asymptomatic manner. However, in cases of immunosuppression, the infection can cause devastating and life-threatening tissue damage, primarily in the brain ^10 5^. The lifelong persistence of JCV and the seemingly high viral load in the human body, as evidenced by the number of viral particles in urine ^11^, contrasts with the low frequency of highly infected cells ^12 13 14 15^. Even under optimal viral infection conditions in vitro, only a small percentage of virus-exposed cells contain significant amounts of virus (Fig. 1a, b). These findings suggest that very few infected cells must produce the virus at high enough levels to ensure consistent output throughout the host’s lifetime.

**Fig. 1:**
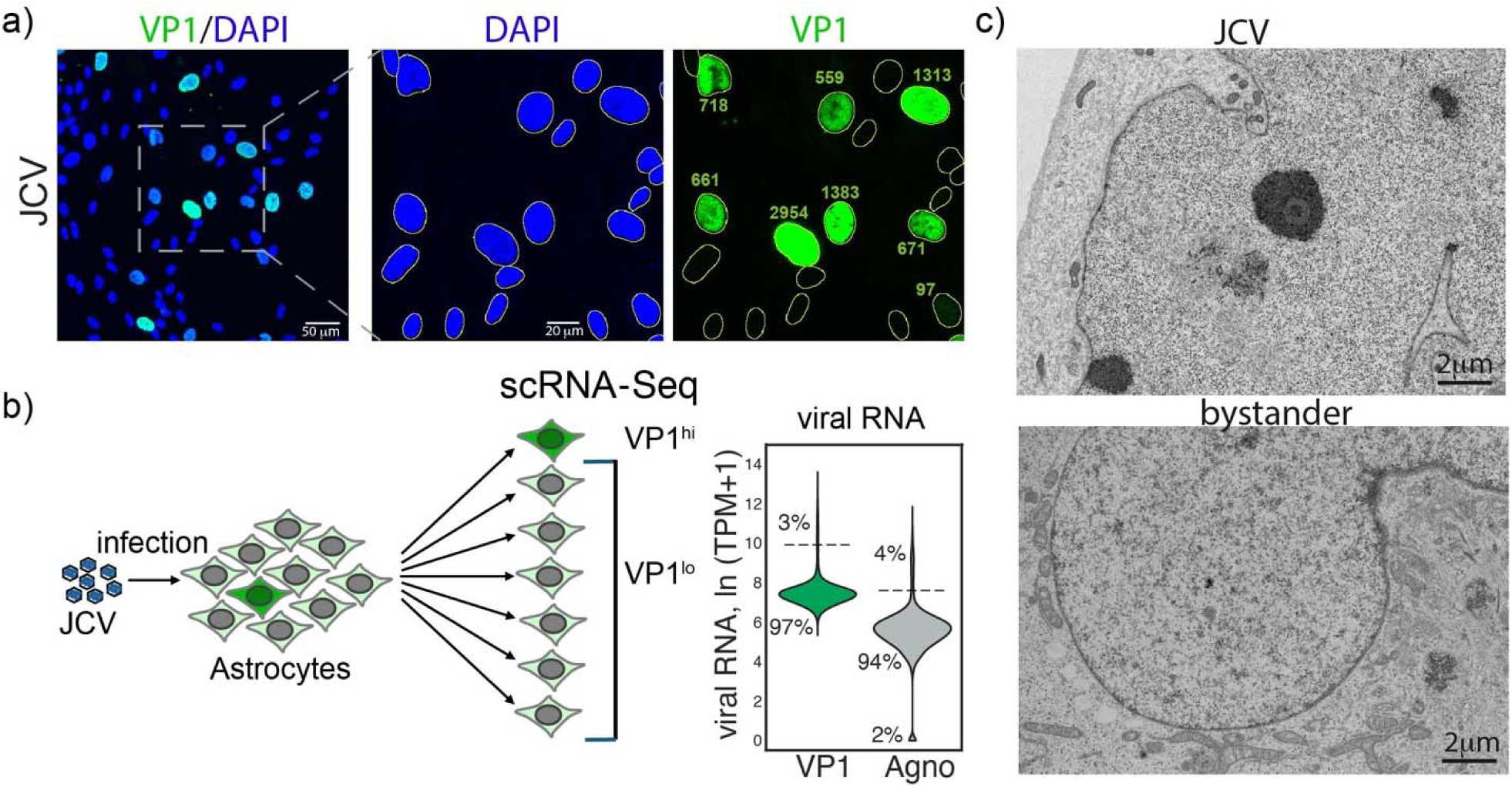
JCV causes productive infection only in a small fraction of astrocytes *in vitro* and accumulates in the nucleus outside high-electron-density heterochromatin areas. a-b) JCV causes productive infection only in a small fraction of astrocytes. a) IF images showing VP1 protein (green; mean signal intensity is shown) and DNA (DAPI; blue). Scale bars are shown in the images. b) Single cell RNA-Seq analysis of JCV infected astrocytes. All infected astrocytes contain viral RNAs (VP1 in green; Agno in grey), but only a small frequency progresses to a VP1^hi^ state. Violin plot shows distribution of expression levels for the individual cells. c) JCV particles accumulate in regions with low chromatin density. EM images show an example of JCV^hi^ cells (top) and JCV bystander cells (bottom). Scale bars are shown in the lower right corner of each image.

We found that infected human astrocyte nuclei contain approximately 1.5 x 10^5^ viral particles. These particles accumulate outside high-electron-density heterochromatin areas, indicating the virus’s ability to generate a space conducive to viral accumulation (Fig. 1c). Insights into the JCV strategy for overcoming constraints of nuclear architecture on viral accumulation come from the analysis of JCV protein sequences. We discovered that one of the JCV proteins, the 8 kDa Agnoprotein (Agno) ^16^, carries motifs similar to those present in nuclear epigenetic regulators that interact with a key chromatin organizer, heterochromatin protein 1 alpha (HP1α) ^17 18^ (Fig. 2a). HP1α utilizes two structurally distinct domains, the chromodomain (CD) and chromoshadow domain (CSD), located at opposite ends of the protein, to bridge di- or tri-methylated histone H3 with several epigenetic regulators and nuclear structure organizers ^17^^19 20 21 22 23.^

**Fig. 2:**
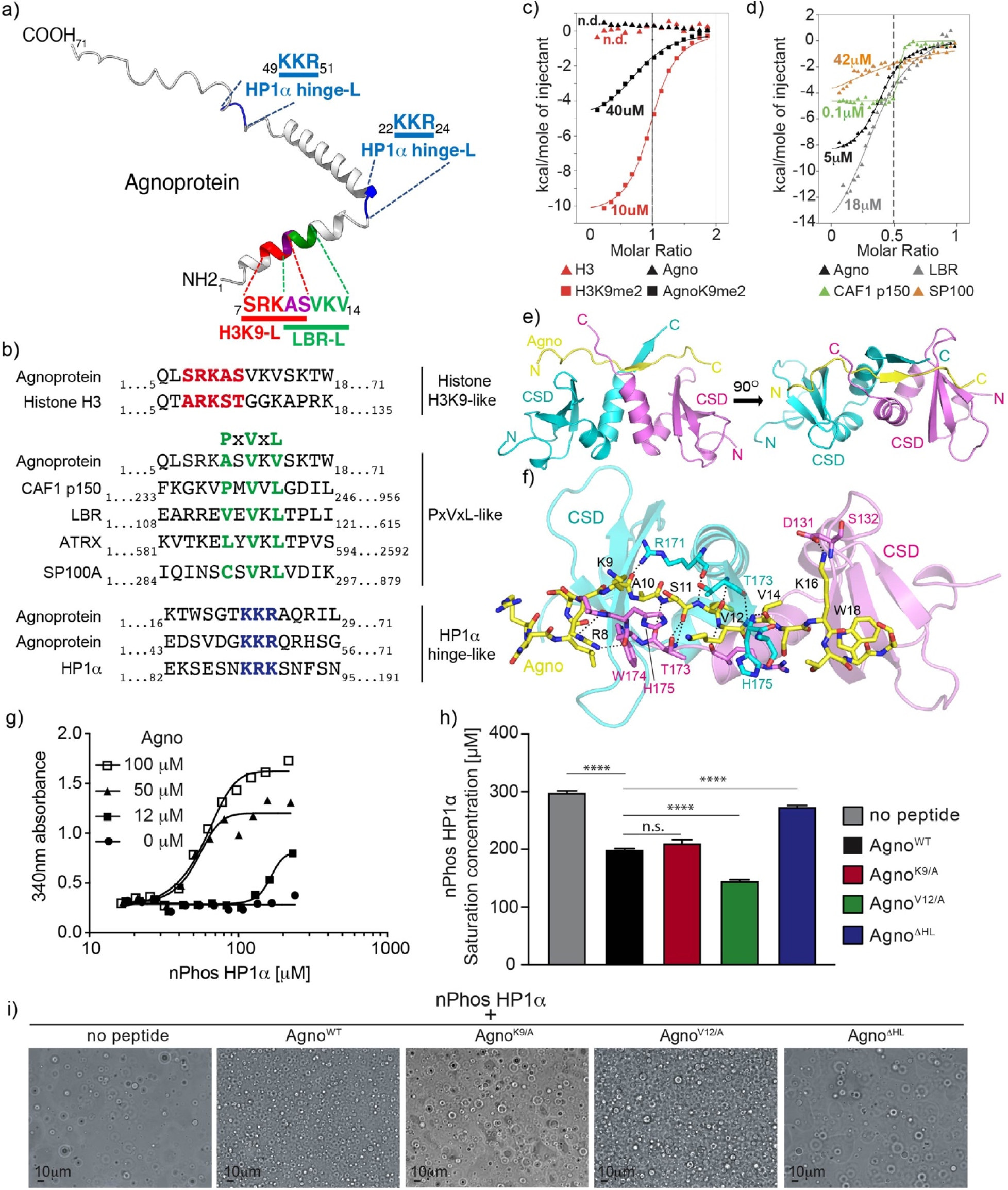
JC virus Agnoprotein contains an array of HP1α-interacting Short Linear Motifs (SLiMs). a) The histone H3K9-like motif (red), the PxVxL-like motif (green) and the HP1α hinge-like motifs (blue) in Agnoprotein are shown. b) The host proteins “mimicked” by Agno. c) The H3K9-like motif in Agno (aa 2-18) binds to the HP1α chromodomain (CD; aa 18-75) in a methylation dependent fashion. d) The PxVxL-like motif of Agno (aa 2-21) binds to the Hp1α chromoshadow domains (CSD; aa 110-191) like other PxVxL motif containing host proteins. c, d) The molecular dissociation constants (K_D_) were quantified by Isothermal Titration Calorimetry (ITC). e, f) The crystal structure of HP1α CSD-Agno complex shows interactions between the HP1α CSD dimer (aa 112-176, C133S/T135S; in green and magenta) and Agnoprotein (aa 3-19; in yellow, red, and blue). Residues involved are denoted and shown in stick representation. g) Agno reduces the saturation concentration necessary for HP1α (nPhos-HP1α) liquid-liquid phase separation. The impact of different concentrations of Agno peptide (aa 2-27) on the saturation concentration of HP1α are shown. Phase separation was measured by turbidity assay at 340 nm. h) The epigenetic mimics in Agno determine the saturation concentration of HP1α required for phase separation as measured by spin-down assay and absorption of the supernatant at 280 nm. P values were determined using ordinary one-way ANOVA (F = 335.6; df = 14). Data show the mean of 3 independent experiments. Error bars represent the SD. i) Micrograph of the liquid-liquid phase separated HP1α samples in the presence of the control and mutant Agno peptides. Scale bar is shown in the lower left corner of each image.

In addition to the relatively large and structured CSD and CD, the hinge region of HP1α connects the CD and CSD and contains a short linear sequence that promotes HP1α’s liquid-liquid phase separation and the formation of HP1α-containing subnuclear chromatin domains ^24^. Remarkably, Agno contains motifs that mimic the CSD and CD interacting sequences in HP1α’s binding partners, as well as the phase separation-inducing motif within the hinge region of HP1α (Fig. 2a, b). Specifically, the SRKAS motif (aa 7-11) of Agno mimics the sequence ARKST (aa 7-11) of histone H3 (Fig. 2b, top), which, upon methylation, can bind to the CD of HP1α ^17^. The adjacent ASVKV sequence (aa 10-14) mimics the PxVxL motif found in numerous nuclear proteins that bind to the CSD of HP1α ^19 20 21 23^ (Fig. 2b, center). Additionally, Agno contains two KKR motifs (aa 22-24 and 49-51) that mimic the phase separation-inducing positive amino acid motif within the HP1α hinge region ^24^ (Fig. 2b, bottom).

These “epigenetic mimics” of Agno can act as bona fide ligands for HP1α. When methylated on lysine 9 (K9) within the H3K9-like motif, Agno can interact with the CD of HP1α with an affinity similar to that of K9-dimethylated histone H3 (Fig. 2c). Agno also directly interacts with the HP1α CSD dimer, similar to other PxVxL motif-containing proteins such as lamin B receptor (LBR), SP100, or CAF1 p150 (Fig. 2d). The high degree of Agno mimicry of host proteins is underscored by atomic details of Agno interaction with HP1α. The crystal structures of the HP1α CSD in complex with Agno, resolved at 3.2 Å, show two CSDs of HP1α forming a dimer, with the two C-termini of the CSD dimer sandwiching Agno to form a triple-stranded β-sheet (Fig. 2e; left shows side view; right shows view from top). This structure is primarily stabilized by intermolecular hydrogen bonds between the backbones of Agno and the CSD C-termini (Fig. 2f). Similar to other reported CSD-interacting proteins ^20 21^, the PxVxL-like motif of Agno embeds into the surface channel of the CSD dimer through hydrophobic interactions (Extended Data Fig. 1a). The essential role of the three hydrophobic amino acid residues of the PxVxL-like sequence in Agno for interaction with the CSD of HP1α is supported by the finding that replacing any of these three amino acids with the negatively charged amino acid glutamic acid (E) completely abolishes detectable interaction (Extended Data Fig. 1b). Similarly, replacing the central valine (V) at position 12 with another hydrophobic amino acid, alanine (A), prevents interaction between the HP1α CSD and Agno, while replacing A10 or V14 with the amino acid A has only minimal impact (Extended Data Fig. 1c).

Agno can also promote the formation of HP1α phase-separated droplets. The addition of Agno peptide to N-terminally phosphorylated HP1α increases phase separation of HP1α in a concentration-dependent manner (Fig. 2g), decreasing the critical concentration required for liquid droplet formation. This positive impact of Agno on HP1α phase separation is strictly dependent on the HP1α hinge-like sequence (HL) in Agno. Mutation of the HL sequence completely abolishes the HP1α phase separation-inducing potential of Agno, while mutations in the other Agno-encoded chromatin mimics have no or opposite effects on HP1α phase separation (Fig. 2h, i).

The ability of Agno to interact with HP1α suggests a potential impact of JCV on chromatin. In astrocytes infected with JCV, DAPI-positive or H3K9me3 heterochromatin accumulates in the center of the cell nuclei, leaving the remaining space potentially available for virus accumulation (Fig. 3a). Centrally located heterochromatin domains contain high levels of HP1α (Fig. 3b). It is conceivable that Agno, which can form dimers or oligomers ^25^, acts as a virus-derived heterochromatin cross-linker and phase separation inducer for chromatin-associated HP1α. Infection of cells with JCV mutant viruses carrying loss-of-function mutations in either the H3K9-like or the PxVxL-like mimic results in a nucleus-wide distribution of HP1α that appears largely dislocated from the DAPI-rich heterochromatin (Fig. 3c).

**Fig. 3:**
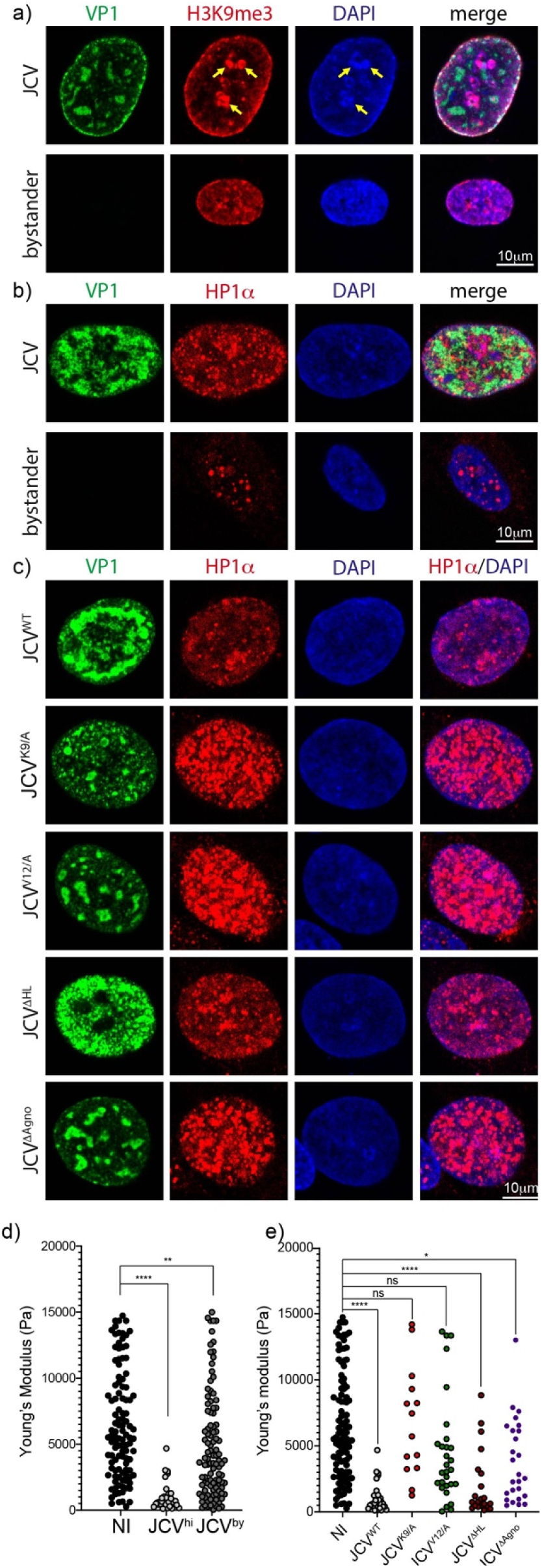
Agnoprotein epigenetic mimics control HP1α nuclear organization and nuclear stiffness. a) JCV promotes formation of heterochromatic domains in the center area of the nucleus (TADs). VP1 (green), H3K9me3 (red) and DNA (DAPI; blue) are shown. The TADs are indicated by yellow arrows. b) HP1α predominantly associates with centrally located heterochromatic domains in JCV^hi^ cells. Immunofluorescence images showing VP1 (green), HP1α (red) and DNA (DAPI; blue). c) HP1α-interacting mimics of Agno play a major role in HP1α organization in the infected nuclei. Representative images of JCV^hi^ cells infected with JCV carrying wild type (WT) Agnoprotein (n = 9), Agnoprotein with inactivating mutations of the epigenetic mimics (n(K9/A) = 9; n(V12/A) = 12; n(ΔHL) = 10), or lacking Agnoprotein (n(ΔAgno) = 12) are shown. VP1 in green, HP1α in red, DNA (DAPI) in blue. a-c) Scale bars are shown in the lower right corner of each image. d) JCV reduces nuclear stiffness. Dot blot showing the Young’s modulus in Pascal (Pa) of individual uninfected astrocytes (n = 110), JCV^hi^ astrocytes (n = 32) or JCV bystander cells (n = 106). The level of infection was quantified by measuring the expression levels of VP1 protein. e) Agnoprotein epigenetic mimics support JCV-induced nuclear softening. Dot plot showing the Young’s modulus in Pa of individual uninfected (n = 110) or JCV^hi^ astrocytes (n = 32) carrying wild type Agnoprotein, Agnoprotein with inactivating mutations of the H3K9-like motif (K9/A; n = 14), the PxVxL-like motif (V12/A; n = 29), the HP1α hinge-like motif (ΔHL; n = 21) or lacking Agnoprotein (ΔAgno; n = 26). d, e) Data are combined from 3 independent experiments. P values were determined using Kruskal-Wallis test. The Young’s modulus values of non-infected and JCV wild type infected astrocytes (JCV^hi^) represented in Fig. 3d and e are identical/from the same measurements.

Heterochromatin plays a defining role in nuclear rigidity ^26 27 28 29^. Therefore, the ability of JCV to interfere with heterochromatin may lead to changes in nuclear rigidity and potentially support virus accumulation by increasing nuclear elasticity. Indeed, we found that JCV infection leads to an almost uniform softening of the nuclei in highly infected (JCV^hi^) cells, but not in bystander cells, compared to control astrocytes (Fig. 3d). The epigenetic mimics in Agno play a key role in Agno-driven nuclear softening in the JCV^hi^ cells. Infection with a virus with inactivating mutations in either the H3K9-like or the PxVxL-like motif in Agno does not cause nuclear softening in JCV^hi^ cells (Fig. 3e). However, mutation of the phase-separation promoting hinge-like sequence in Agno did not impact the Agno-driven nuclear softening observed for the wild-type JC virus (Fig. 3e). It is likely that the chromatin-dislocated HP1α-containing nuclear speckles that we observed upon infection with viruses carrying inactivating mutations in either the CD- or CSD-interacting motifs (Fig. 3c) can support nuclear stiffness and therefore prevent JCV-induced nuclear softening.

The key role of Agno in JCV-mediated nuclear softening is in stark contrast to the lack of Agno’s effect on gene expression in the infected cells. Using single-cell RNA analysis, we identified 130 and 56 genes that become upregulated or downregulated, respectively, in the JCV^hi^ cells (Fig. 4a). Despite the effect on heterochromatin, JCV has no effect on the expression of the heterochromatin-silenced genes, such as endogenous retroviruses or transposable elements ^30 31^ (Extended Data Fig. 2a). The majority of the upregulated genes in the JCV^hi^ cells belong to groups involved in DNA repair, DNA replication, biogenesis, and cell cycle regulation (Extended Data Fig. 2b)— processes essential for JCV DNA replication and primarily driven by the expression of viral large T-antigen ^32 33 34^. Most importantly, the observed changes in gene expression did not depend on the presence of Agno (Fig. 4b).

**Fig. 4:**
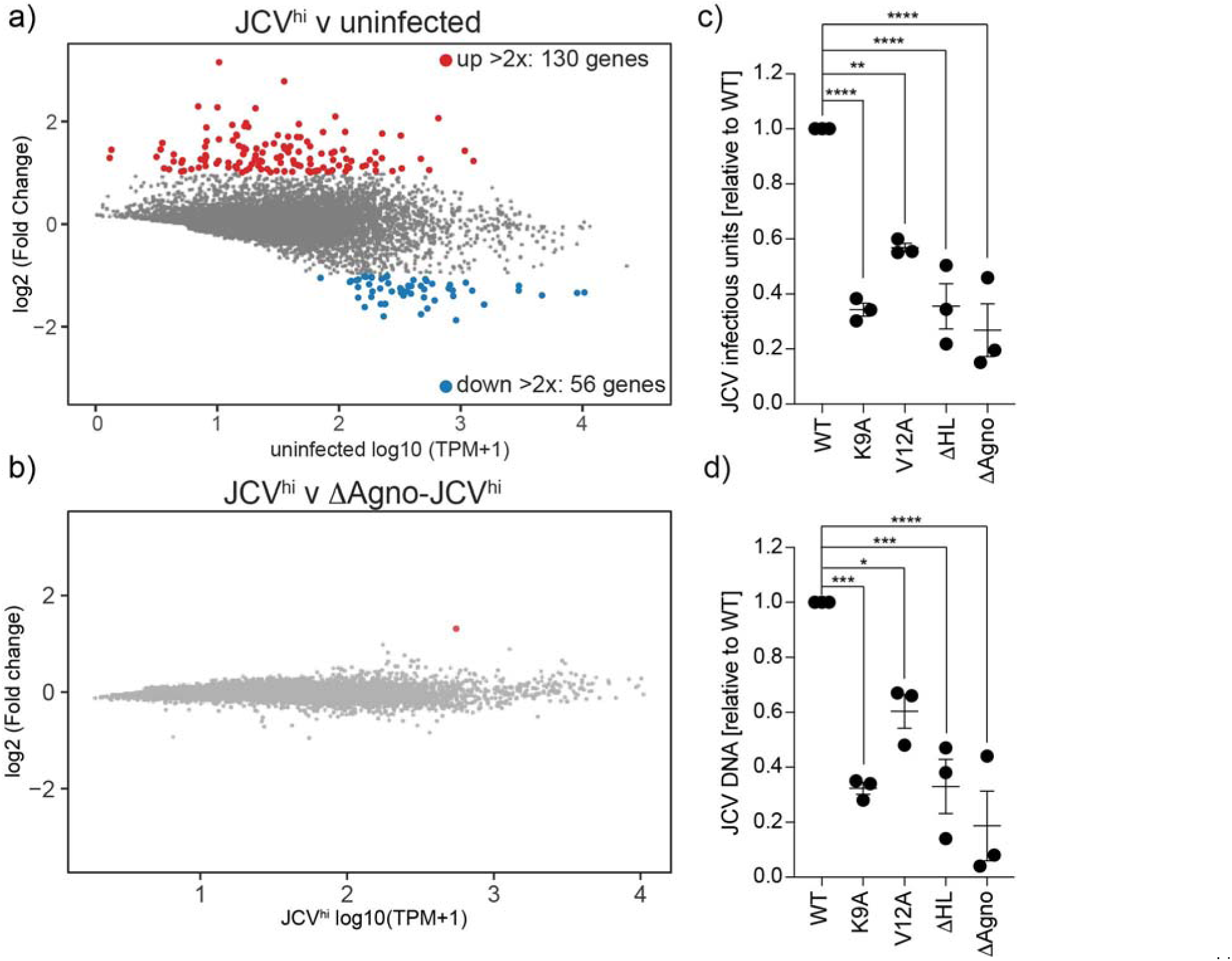
Agnoprotein epigenetic mimics control viral infection without impacting host gene expression. a) JCV infection affects the expression of a limited number of host genes in the JCV^hi^ astrocytes. The gene expression differences have been measured by single cell RNA sequencing. b) Agno does not contribute to host gene expression changes in JCV^hi^ cells. Differential gene expression analysis of JCV^hi^ cells infected with wild type or Agno-deficient JCV. c) The infectious titers of cell lysates infected with wild type JCV and JCV epigenome mimic mutants 28 days after viral genome transfection are shown. d) Relative JCV genome levels in the lysates of non-infected or infected cells (wild type JCV and JCV epigenome mimic mutants) are shown. c, d) Shown is the mean with SEM of 3 independent experiments. P values were determined using ordinary one-way ANOVA (for d: F= 26.25; df = 14; for e: F = 17.33; df = 14).

The impact of JCV on chromatin and the mechanical properties of cell nuclei plays an important role in viral replication. Glia-like SVG-A cells ^35 36^ were transfected with equal amounts of control or mutant viral genomic DNA containing inactivating mutations of the epigenetic mimics, followed by quantification of infectious virus production. The mutations in Agno epigenetic mimics reduced JCV virus accumulation (Fig. 4c), confirmed by a similarly strong decrease in viral genome copies in the infected cell lysates (Fig. 4d).

In summary, our data provide a paradigm-shifting view of the mechanism of JC virus-host interaction, where the virus’s impact on nuclear mechanics appears to promote virus-host cohabitation rather than perpetual conflict involving the costly maintenance of antiviral response programs. The remarkable use of epigenetic mimicry by the virus to impact nuclear mechanics—which we define as a supra-epigenomic impact—and the key role of these mimics in viral infection suggest the critical role of JCV epigenetic mimicry in viral maintenance and expansion, especially during JCV-induced human diseases, including brain inflammation and potentially cancer. It is likely that other chronically persisting human or animal DNA viruses that replicate and form in the nucleus also possess the ability to interfere with nuclear mechanics in a JC virus-like fashion, using similar or different types of epigenetic mimics embedded within their proteins.

## Methods

### Cell culture

SVG-A cells^35,36^ (kindly provided by Dr. Walter J. Atwood) were cultured in Eagle’s Minimum Essential Medium (EMEM; ATCC; 30-2003) supplemented with 10% FBS (Corning; 35-010-CV; Lot 12522001), 100μg/ml Penicillin/Streptomycin and 2mM L-Glutamine.

Primary human astrocytes (Creative Biolabs; NCC20-9PZ01; Lot NR34) were cultured in AGMTM Astrocyte Growth Medium from Lonza (BulletKit CC-3186) and used between passage numbers 2 to 4.

All cells were grown at 37°C in a 5% CO_2_ atmosphere in a Heraeus Kendro HeraCell 150 CO_2_ Incubator.

### Generation of mutant JCV genomes

The plasmid to generate JC virus strain Mad1-SVEΔ (pUC19-Mad1-SVEΔ), a subclone of the Mad1-SVEΔ hybrid strain of JCV^37^, was kindly provided by Dr. Walter Atwood. Inactivating mutations of the HP1α-interacting chromatin protein mimicry in JC virus Agnoprotein were introduced using the QuikChange II XL Site-Directed Mutagenesis Kit from Agilent (#200521) and the appropriate primers (Sigma; sequences see below) following the manufacturer’s instructions. All mutations in Agnoprotein were confirmed by Sanger DNA sequencing (Macrogen).

Agno_K9/A_forward: CGCCAGCTGTCACGTGCTGCTTCTGTGAAAGTTAGTAAAACCTGG
Agno_K9/A_reverse: CCAGGTTTTACTAACTTTCACAGAAGCAGCACGTGACAGCTGGCG
Agno_V12/A_forward: GCTGTCACGTAAGGCTTCTGCTAAAGTTAGTAAAACCTGGAGTGG
Agno_V12/A_reverse: CCACTCCAGGTTTTACTAACTTTAGCAGAAGCCTTACGTGACAGC
Agno_ΔHL_forward (K22/A, K23/A, R24/A): CTGTGAAAGTTAGTAAAACCTGGAGTGGAACTGCAGCAGCAGCTCAAAGGATTTTAATTTTTTTGTTAGAATTTTTGCTGG
Agno_ΔHL_reverse (K22/A, K23/A, R24/A): CCAGCAAAAATTCTAACAAAAAAATTAAAATCCTTTGAGCTGCTGCTGCAGTTCCACTCCAGGTTTTACTAACTTTCACAG
ΔAgno_forward: GCTTGTCACCAGCTGGCCATAGTTCTTCGCCAGCTGTCAC
ΔAgno_reverse: GTGACAGCTGGCGAAGAACTATGGCCAGCTGGTGACAAGC

### Generation of wild type and mutant JC virus stocks

The viral genome was digested by the restriction enzyme BamHI-HF for 2 hours at 37°C. The digested DNA was precipitated via Acetate/Ethanol precipitation. The efficiency of the digest of the plasmid was checked on a 1% agarose gel. 2×10^6^ SVG-A cells were plated in a T-75 flask and allowed to adhere overnight. 11ug of linearized DNA containing the viral genome was transfected into SVG-A cells using Lipofectamine 3000 following the manufacturer’s instructions. The medium was changed 12 hours (h) after transfection. Once cells reached 95% confluency, the transfected cells were transferred into a T-175 flasks and cultured for 28 days (onset of the virus-induced cytopathic effect). During this time the culture medium was replaced every 4 days. The cells were detached by cell scraping, pelleted by centrifugation, and then resuspended in 3ml 20mM Tris-HCl (pH 7.4) containing 0.2% FBS. To release JC virus from the producing cells, the cells underwent 3 freeze/thaw cycles followed by treatment with 0.05U/ml neuraminidase type V (Sigma-Aldrich; N2876) at 37°C for 16 hours. The neuraminidase was heat inactivated at 56°C for 30 minutes. Cell fragments were removed by centrifugation at 10000xg for 20 minutes. The supernatant was aliquoted, and aliquots were stored at −80°C until usage.

### Titration of JCV infectious units

Viral titers were determined by incubation of SVG-A cells with serial dilutions of the virus containing cell lysates for 12h, followed by culturing the cells for 6 days. At day 6, cells were collected, fixed and permeabilized using the eBioscience™ Foxp3 / Transcription Factor Staining Buffer Set (Invitrogen; 00-5523-00) following the manufacturer’s instructions. The cells were stained with primary antibodies targeting the viral capsid protein VP1 (Creative Biolabs; MOB-279YC; Lot# CB2210Y36; 1:100 dilution) and Agnoprotein (custom order generated by Covance Research Products Inc. in Denver against the following peptide: CGKKRQRHSGLTEQTYSALPEPKAT; 1:400 dilution) for 30 minutes at 4°C followed by secondary antibody staining (donkey anti-mouse IgG Alexa Fluor 568, Fisher Scientific Co., # A10037; donkey anti-rabbit IgG Alexa Fluor 647, Fisher Scientific Co., #A31573 or donkey anti-rabbit IgG Alexa Fluor 488, Fisher Scientific Co., #A21206; 1:200 dilution) for 30 minutes at 4°C with 2 washing steps for 5 minutes at room temperature with FACS permeabilization buffer after each antibody staining. The stained cells were acquired using a BD LSRFortessa Cell Analyzer and BD FACSDiva software version 8.0. Data were analyzed using FlowJo10 software. The frequencies of cells expressing JC virus VP1 were determined by comparison to non-infected cells, and the number of infectious units per ml of viral lysate were calculated. The results of 3 independent experiments were plotted in GraphPad Prism version 8. P values were determined using ordinary one-way Anova. P>0.05, ns; p<0.05, *; p<0.01, **, p<0.001, ***; p<0.0001, ****.

### PCR for viral genome quantification

To determine viral genome copies in the virus containing cell lysates, DNA from 30μl cell lysates were isolated using the E.Z.N.A. Tissue DNA Kit (omega BIO-TEK; #D3396-01) following the manufacturer’s instructions. The purified DNA was 1:10 diluted with dH_2_O and then used for DNA copy quantification via qPCR using LightCycler 480 SYBR Green I Master (Roche, #04887352001) and the following primers on a Roche LightCycler 480 Real-Time PCR system:

JCVP1-45F: ggtgacaacttatacttgtcagctgtt
JCVP1-812R: tgctgggaaccagacctgtt

The results of 3 independent experiments were plotted in GraphPad Prism version 8. P values were determined using ordinary one-way Anova. P>0.05, ns; p<0.05, *; p<0.01, **, p<0.001, ***; p<0.0001, ****.

### Electron microscopy of JC virus infected astrocytes

Astrocytes were infected by incubation with JCV diluted in astrocyte growth medium at an MOI of 0.6 for 12h. The medium was changed every 3 days. 12 days after infection the cells were fixed in 2% glutaraldehyde in 0.08 M sodium cacodylate buffer (pH 7.2) containing 2 mM CaCl_2_ for 1h at room temperature and postfixed in 1% osmium/0.8% potassium ferricyanide in 0.1 M cacodylate buffer, followed by post-staining in 1% uranyl acetate in water, dehydration in an ethanol series, and embedding in Eponate 12 (Ted Pella, Inc). Ultrathin sections (60–65 nm) were stained with uranyl acetate and lead citrate, and images were acquired using a Tecnai 12 Spirit transmission electron microscope (FEI, Hillsboro, Oregon) operated at 120 kV, equipped with an AMT BioSprint29 digital camera.

### Immunofluorescence imaging

Astrocytes, cultured on 35mm imaging dishes with ibidi Polymer Coverslip bottom with low walls and an imprinted 500 μm cell location grid (Ibidi; # 80156), were left uninfected or were infected with JCV for 12 days as described above. At day 12 post infection, cells were fixed with 4% paraformaldehyde (PFA) in PBS for 15 minutes at room temperature (RT). After 2 washes with PBS for 10 minutes, cell membranes were permeabilized in IF permeabilization buffer (0.1% sodium citrate and 0.1% Triton X-100 in dH_2_O) for 5 minutes, followed by two washes with PBS for 5 minutes. After blocking in IF blocking buffer (2.5% BSA, 0.1% Tween 20 and 10% normal donkey serum in PBS) for 1h at RT, cells were strained with the corresponding primary antibodies overnight (ON) at 4°C (mouse anti-VP1, Creative Biolabs, MOB-279YC, Lot# CB2210Y36; 1:100 dilution; rabbit anti-Agnoprotein, Covance Research Products Inc., 1:300 dilution; rabbit anti-H3K9me3; Active Motif, #39285, 1:300; rabbit anti-HP1α, Cell Signaling, #2616S, 1:200 dilution; rabbit anti-Lamin A/C, Abcam, ab133256, 1:300 dilution). Cells were washed 3 times for 15 minutes at RT with IF washing buffer (0.25% BSA and 0.1% Tween 20 in PBS), then stained for 1h at RT with the corresponding fluorophore-coupled secondary antibodies (Dn anti-Mouse IgG Alexa Fluor 488, Fisher Scientific CO., # A21202; Dn anti-Rabbit IgG Alexa Fluor 568, Fisher Scientific CO., # A10042). After secondary antibody staining, cells were stained for 20 minutes with NucBlue Fixed Cell Stain ReadyProbes reagent (Fisher Scientific Co.; R37605) in PBS according to the manufacturer’s instructions. Cells were washed 2 more times with IF washing buffer and stored in PBS with 0.01% NaN_3_ until image acquisition.

Z-stack images were acquired on a Zeiss LSM 880, AxioObserver microscope with a 63x/1.40 Oil objective in the Rockefeller University’s Bio-Imaging Resource Center, RRID:SCR_017791.

### Force indentation spectroscopy of JC virus infected NHA cell nuclei using AFM

Atomic Force Microscopy (AFM) mechanical measurements were performed on noninfected and JCV infected astrocytes 12 days after infection (as described above), cultured on 35mm imaging dishes with ibidi Polymer Coverslip bottom with low walls and an imprinted 500 μm cell location grid (Ibidi; # 80156). The AFM measurements were performed using a JPK NanoWizard 4 (Bruker Nano) atomic force microscope mounted on an Eclipse Ti2 inverted fluorescent microscope (Nikon) and operated via JPK SPM Control Software v.6. MLCT triangular silicon nitride cantilevers (Bruker) were used to access nuclei stiffness. Forces of up to 3 nN were applied at 10 micron per second constant cantilever velocity and ensuring an indentation depth of 500 nm. All analyses were performed with JPK Data Processing Software v.6 (Bruker Nano) by first removing the offset from the baseline of raw force curves, then identifying the contact point and subtracting cantilever bending before fitting the Hertz model with correct tip geometry to quantitate the Young’s Modulus. The use of gridded plates allowed identification of infection state of mechanically characterized NHA post AFM measurements by immunofluorescence analyses of the intracellular VP1 signal (immunofluorescence staining of VP1 and DNA as described above) on a Nikon eclipse Ti2 inverted microscope mounted with a CSU-W1 spinning disk (Nikon). The results of 3 independent experiments were plotted in GraphPad Prism version 8. P values were determined using ordinary one-way Anova. P>0.05, ns; p<0.05, *; p<0.01, **, p<0.001, ***; p<0.0001, ****.

### Protein expression and purification for in vitro binding studies via ITC

The sequences corresponding to HP1α CD (amino acid (aa) 18-75) or CSD (aa 110-191) were cloned into a pRSFDuet-1 vector (Sigma-Aldrich; #71341 (Novagen)) engineered with an N-terminal hexa-histidine (His_6_)-SUMO tag. The fusion proteins were expressed in *E. coli* strain BL21 (DE3) RIL (Stratagene, Santa Clara, CA, USA). The cells were cultured at 37°C until the OD_600_ reached 0.8. The media was cooled to 18°C and IPTG was added to a final concentration of 0.4 mM to induce protein expression overnight at 16°C. The cells were harvested by centrifugation at 4°C and disrupted by sonication in buffer W (500 mM NaCl, 20 mM imidazole, and 50 mM Tris-HCl, pH 7.5) supplemented with 1 mM phenylmethylsulfonyl fluoride (PMSF) protease inhibitor and 2 mM β-mercaptoethanol. After centrifugation, the supernatant was loaded onto a HisTrap FF column (GE Healthcare, Little Chalfont, Buckinghamshire, UK). After extensive washing with buffer W, the target protein was eluted with buffer W supplemented with 300 mM imidazole. The hexa-histidine-Sumo tag was cleaved by Ulp1 protease and removed by two times elution through a HisTrap FF column. Individual pooled target proteins were further purified by passage through a Hiload 16/60 Superdex 200 column (GE Healthcare) with buffer E (150 mM NaCl, 10 mM Tris-HCl, pH 7.5, and 1 mM DTT).

### ITC measurements

All binding experiments were performed on a Microcal ITC 200 calorimeter. Purified HP1α constructs were dialyzed overnight against buffer A (150 mM NaCl, 1 mM DTT, and 10 mM Tris, pH 7.5) at 20°C. Lyophilized Agnoprotein and analogues peptides were purchased from the Rockefeller University Proteomics Core Facility and dissolved in buffer A. The assays were performed with 1 mM peptides and 0.2 mM HP1α proteins. The exothermic heat of the reaction was measured by sequential injections of the peptide into the protein solution, spaced at intervals of 180 s. The titration was done according to the standard protocol at 20°C and the data were fitted using the program Origin 7.0 with one site model.

Peptide sequences for ITC measurements with CD:
Agno (aa 2-18): VLRQLSRKASVKVSKTW
AgnoK9me2 (aa 2-18): VLRQLSRK_me2_ASVKVSKTW
H3 (aa 1-17): ARTKQTARKSTGGKAPRY
H3K9me2 (aa 1-17): ARTKQTARK_me2_STGGKAPRY

Peptide sequences for ITC measurements with CSD:
Agno (aa 2-21): VLRQLSRKASVKVSKTWSGT
hCaf1 p150 (aa 231-249): ILFKGKVPMVVLQDILAVR
hLBR (aa 103-122 plus N-terminal Y): YQADIKEARREVEVKLTPLIL
hSP100 (aa 279-298 plus C-terminal Y): LHNHGIQINSCSVRLVDIKKY
Agno A10/L (aa 2-21): VLRQLSRKLSVKVSKTWSGT
Agno A10/E (aa 2-21): VLRQLSRKESVKVSKTWSGT
Agno V12/A (aa 2-21): VLRQLSRKASAKVSKTWSGT
Agno V12/E (aa 2-21): VLRQLSRKASEKVSKTWSGT
Agno V14/A (aa 2-21): VLRQLSRKASVKASKTWSGT
Agno V14/E (aa 2-21): VLRQLSRKASVKESKTWSGT

### Crystallization and structure determination

The sequences corresponding to the HP1α CSD (aa 112-176) was cloned into a pRSFDuet-1 vector (Novagen) engineered with an N-terminal hexa-histidine (His_6_)-SUMO tag. To improve the crystallization, the dual mutations C133S/T135S was introduced to the HP1α CSD (aa 112-176) using the QuikChange Mutagenesis Kit (Stratagene). HP1α was expressed and purified as described above. 5-FAM-labelled JCV Agnoprotein (aa 3-19) was chemically synthesized by the Rockefeller Proteomics Core Facility and dissolved in buffer E to 5 mg/mL. To assemble the CSD-Agnoprotein binary complex, the purified HP1α CSD (aa 112-176, C133S/T135S) protein was mixed with 5-FAM-labelled JCV Agnoprotein (aa 3-19) at the molar ratio of 1:5 in buffer C (300 mM NaCl, 20 mM Tris-HCl, pH 7.5, and 1 mM DTT), and incubated on ice for 1h. Crystallization conditions were determined with crystal screens (Qiagen) by sitting-drop vapor diffusion with a Mosquito crystallization robot (TTP Labtech). Crystallization conditions were optimized using the hanging drop vapor diffusion method at 20°C. Crystals were grown from drops consisting of 1 μL protein solution (about 10 mg/ml) and 1 μL reservoir solution containing 0.01 M NiCl_2_, 0.1 M Tris-HCl, pH 8.5, and 20% PEG2000MME (v/v).

Peptide sequence for crystallization with CSD: 5-FAM Agno (aa 3-19): FAM-LRQLSRKASVKVSKTWS

The x-ray diffraction data set was collected on the 24-IDE beamline at the Advanced Photon Source (APS) at 100K at the Argonne National Laboratory. The wavelength was 0.9792Å. The diffraction data were indexed, integrated, and scaled using the HKL2000 package^38^, and the structure of HP1α CSD-Agnoprotein complex was solved using molecular replacement with PHENIX^39^, using a modified HP1β CSD-CAF1 structure (PDB code: 1S4Z) as the search model. Model building and structural refinement was carried out using COOT^40^, O^41^ and PHENIX. The statistics of the data collection and refinement are shown in the Supplementary Table-1. Figures were generated using PyMOL (http://www.pymol.org). The structural coordinates have been deposited into the Protein Data Bank (PDB) under the accession number 9CMT.

### Turbidity based determination of the HP1**α** phase separation behavior

Serial dilutions of purified nPhos HP1α (protein purification as described in Larson et al., Nature, 2017^24^) were performed in 96 well liquid reservoirs. 19μL of these samples were added to a clear bottom 384 well plate (Corning) and absorbance was read at 340 nm in a Spectramax M5 plate reader at 25°C. To address the impact of Agno peptides on nPhos HP1α phase separation, 1 μL of peptide at the appropriate concentration was added to the HP1α solutions, mixed, and incubated for 5 minutes before reading absorbance at 340nm. The final Agno peptide concentrations are indicated in the figure.

### Approximation of the critical concentration by spin down method

A ∼300µM nPhos HP1α solution was incubated with various Agno peptides at a final concentration of 100µM in a total volume of 10µl. The samples were incubated for 5 minutes at 25°C, then centrifuged at 10,000xg for 5 minutes in a microcentrifuge. The concentration of HP1α in the supernatant, a proxy of the critical concentration of HP1α to undergo phase separation, was determined with a nanodrop instrument by A280 absorbance in triplicate. All measurements were plotted, and statistical tests were performed using GraphPad Prism version 8. P>0.05, ns; p<0.05, *; p<0.01, **, p<0.001, ***; p<0.0001, ****.

### Imaging of nPhos HP1**α** liquid droplets

Samples were prepared as described for the spin down method. 5µl of the sample were loaded onto a flow cell made from a cover slide and a peg-silane treated coverslip. 1 minute after loading the samples, micrographs were collected on a Zeiss Axiovert 200M microscope using a 20X phase contrast objective.

Peptide sequences for HP1α liquid-liquid phase separation experiments:
Agno (aa 2-27): VLRQLSRKASVKVSKTWSGTKKRAQR
Agno_K9/A (aa 2-27): VLRQLSRAASVKVSKTWSGTKKRAQR
Agno_V12/A (aa 2-27): VLRQLSRKASAKVSKTWSGTKKRAQR
Agno_ΔHL (K22/A, K23/A, R24/A; aa 2-27): VLRQLSRKASVKVSKTWSGTAAAAQR

### scRNA-Sequencing using 10xGENOMICS

Astrocytes were plated in 6-wells (Corning; #3516) and were either left uninfected or were infected with JCV as described above. 10 days after infection, astrocytes were collected by trypsinization (LONZA WALKERSVILLE INC.; CC-5034), filtered through a Falcon 70μm cell strainer (Corning; 352350) and then directly used for single cell RNA-Seq library generation using the Chromium Single Cell 3’ GEM, Library & Gel Bead Kit v3 (16 rxns; PN-1000075) and a 10x Chromium controller according to manufacturer’s instructions. The libraries were sequenced on an Illumina NovaSeq 6000 platform using the NovaSeq *6000* v1.5 Reagent Kit with the following sequencing parameters: 28X10X10X90.

### Single-cell transcripts mapping

The masked genome reference GRCh38, and the corresponding gene transfer format (gtf) annotation file for the gene reference were downloaded from ftp://ftp.ensembl.org/pub/release-96/gtf/homo_sapiens. The downloaded GRCh38 reference was then combined with JC virus sequences and transposable elements (TE) sequences to create a custom reference and annotation files for the data alignment. Briefly, the JC virus sequences were divided into separate regions corresponding to Agno (101-316 bp), VP2 and VP3 overlapping gens (350-1384bp), and VP1 gene (1385-2357bp)^42^ and added to the fastq and gtf files. The sequences for small (273bp) and large T antigen (1821bp) were also added to the reference file. Finally, in order to include TEs in the analysis, 427 consensus TE fastq sequences from Repbase^43^ representing LINE, SINE, and LTRs TE families that are sufficiently distinct in terms of their repeats to allow for unique read mapping were downloaded. These TE sequences as well as the annotation gtf file that we have created were merged with above files to generate the final reference and annotation files. The final custom reference was then built and the reads were mapped to human and viral genes, and TE subfamilies using the Chromium 10x workflow and the Chromium 10x Cell Ranger Single-Cell Software Suite (http://software.10xgenomics.com/single-cell). Only the unique alignments were considered and counted for the differential expression of TE analysis. The number of differences is defined as the number of differences in TE subfamilies. To estimate RNA velocities for our single cell data we used velocyto run10x from the Velocyto package on the Chromium 10x Cell Ranger output files as suggested by the velocyto workflow^44^.

Data preprocessing and clustering analysis was done with Scanpy v.1.6.0. First, cells with low UMI counts (<=2000) were removed. Next, cells were filtered by removing high UMI cells (*i.e.,* log10 transformed UMIs > mean + 2 standard deviations in each dataset). This step removed cells with total UMI counts > 55423 for the astrocyte dataset. Finally, cells with a high fraction of reads (>=20%) mapping to mitochondrial genes were removed.

For clustering analysis, host genes detected in at least three cells were retained. The digital gene-expression matrix was first normalized by the total UMI count per cell and log-transformed after adding a pseudo count. Next, the top 5,000 genes with the highest variance were selected, with batch-effect regressed out by the total UMI count. The gene count matrix was then scaled to unit variance. The dimensionality of the data was reduced by principal component analysis (PCA) (50 components) first and then with UMAP, followed by Leiden clustering performed on the top 50 principal components (resolution = 1.5) by scanpy v.1.6.0. For UMAP visualization, the PCA matrix was directly fit into the scanpy.tl.umap function with min_dist of 0.3 and default settings. After the preprocessing and clustering steps mentioned above, four clusters corresponding to apoptotic or damaged cells (median UMI count of 4,246.5 vs. 17,379 of the other cells) were filtered out before downstream analysis.

For every cell, the viral RNA expression level was quantified as the number of viral transcripts per million host UMIs (TPM). This value was further log-transformed after adding a pseudo count. We chose a cutoff value distinguishing highJCV (VP1^hi^ or JCV^hi^) and lowJCV (VP1^lo^ or JCV^by^) cells based on the expression distribution of the VP1 gene: cells with log-TPM (VP1) higher than ten were tagged as VP1^hi^/JCV^hi^ cells, and the rest were counted as VP1^lo^/JCV^by^ cells.

Differential gene expression (DE) analysis comparing JCV^hi^ or JCV^by^ cells to control cells was done with Seurat v4.0.3. Raw data were first normalized using the SCTransform function. DE genes between different groups were identified using the FindMarkers function. Finally, the identified DE genes were filtered by an adjusted p-value cutoff of 0.05.

Raw and processed data were deposited at the NCBI Gene Expression Omnibus (GEO accession number: GSE271609, reviewer’s token: unezeyeitnghxon).

### Statistical analysis

All experiments were conducted with an appropriate sample size that was estimated based on our previous experience. No statistical methods were used to predetermine sample size. Dot plots were routinely used to show individual data points for each experimental observation. Statistical analyses were performed using GraphPad Prism 8. Normality and Lognormality tests were performed to determine if ordinary one-way ANOVA (normal distribution) or Kruskal-Wallis test (no normal distribution) is used for statistical analysis. *P* values <0.05 were considered significant; \**P* < 0.05, \*\**P* < 0.01, \*\*\**P* < 0.001, \*\*\*\**P* < 0.0001. *P* values >0.05 were considered non-significant (ns). The sample sizes and statistical tests used are stated in each figure legend. For ANOVA, the F value (F) and degree of freedom (df) are stated in each figure legend.

## Data availability

Raw and processed single cell gene expression data were deposited into the NCBI Gene Expression Omnibus database under GEO accession number GSE271609 (reviewer’s token: unezeyeitnghxon) and are available at the following URL: https://www.ncbi.nlm.nih.gov/geo/query/acc.cgi

The structural coordinates have been deposited into the Protein Data Bank (PDB) under the accession number 9CMT.

All other primary data that support this study are available from the corresponding authors upon reasonable request.

## Code availability

All custom codes and scripts used for single cell gene expression analysis in this study are available from GitHub at the following URL: https://github.com/mb3188/JC_virus_reference_and_gtf

## Acknowledgements

We thank Hilda Amalia Pasolli from the Rockefeller University’s Electron Microscopy Resource Center for help with EM sample preparation and image acquisition, Henry Zebroski, III, from The Rockefeller University Proteomics Resource Center (<u>RRID:SCR_017797</u>) for synthesis of all peptides used in this study, and the staff of the Rockefeller University’s Genomics Resource Center (RRID:SCR_020986) for scRNA-Seq library generation and sequencing. We thank Walter J. Atwood (Brown University) for providing SVG-A cells and the JC virus strain Mad1-SVEΔ genome-containing plasmid pUC19-Mad1-SVEΔ to us. We thank Puja Majumder for processing the ITC data. This work was supported by the Stavros Niarchos Foundation supporting Global Infectious Disease Research at the Rockefeller University awarded to A.T., the Intramural Research Program of the National Institute of Diabetes and Digestive and Kidney Diseases (NIDDK), National Institute of Health (NIH) awarded to Y.A.M., the National Natural Science Foundation of China Grant 82151216 awarded to W.X., the Maloris Foundation and the Memorial Sloan-Kettering Cancer Center Core Grant (P30CA008748) in support of D.J.P. laboratory, the NIH R35 GM127020 grant awarded to G.J.N., and the Rockefeller Start-up funding awarded to J.C.. Z.L. is supported by the David Rockefeller Graduate Program in Bioscience at The Rockefeller University, and V.S. is supported by the Beckman Foundation award ID: AWD00000407 for Light-Sheet microscopy and Data Science.

## Contributions

U.S. and A.T. conceived and designed the project. U.S. designed and executed most of the experiments. Y.A.M. performed and analyzed force indentation experiments to address the impact of virus on nuclear stiffness. Y.A.M. and S.A.W. contributed to the design of the force indentation experiments. W.X. generated the crystal of Agno-HP1α CSD complex. W.X., K.C. and Y.G. performed structural analysis. S.C. performed and analyzed ITC measurements. D.J.P. supervised ITC experiments and protein structure determination. V.P.S. contributed to imaging data analysis. A.G.L. performed liquid-liquid phase separation experiments and analysis. G.J.N. supervised liquid-liquid phase separation experiments. Z.L. and M.B. performed single cell gene expression analysis. J.C. supervised single cell gene expression analysis. Y.A.M., S.A.W., G.J.N. and D.J.P. contributed to data interpretation. A.T. supervised the entire project. A.T. and U.S. wrote the manuscript with input from all coauthors.

## Ethics declarations

**Extended data Fig. 1:**
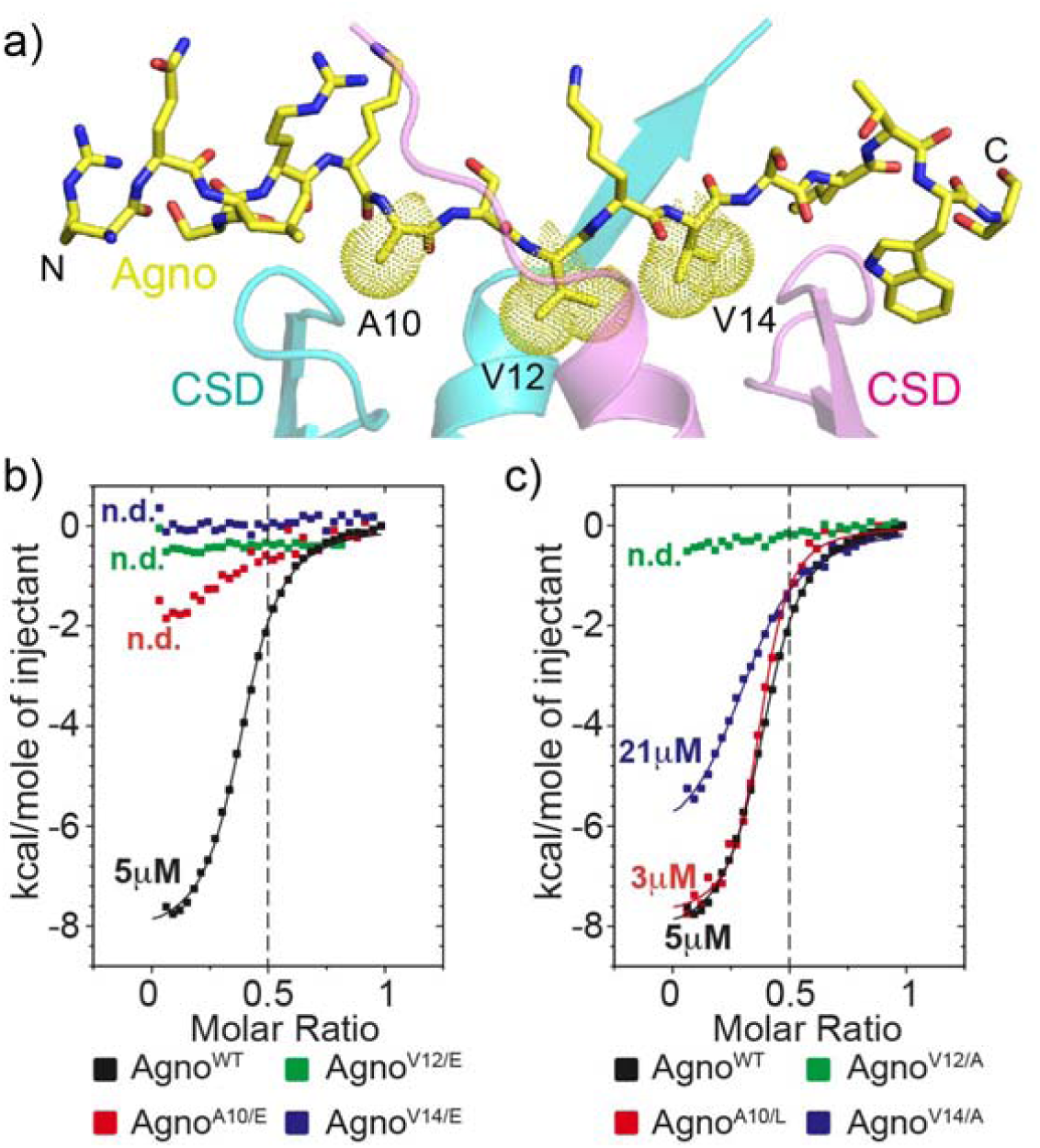
JCV Agno binds to the CSD of HP1α via a PxVxL-like sequence. a) The crystal structure of HP1α CSD-Agno complex shows interactions between the HP1α CSD dimer (in green and magenta) and Agnoprotein (in yellow, red, and blue). Residues involved are denoted and shown in stick representation. b) The three hydrophobic residues of the PxVxL-like motif in Agno (aa 2-21) are essential for the interaction with the Hp1α chromoshadow domain (aa 110-191). c) V_12_ of Agno plays a central role in the interaction with the HP1α CSD. b, c) The molecular dissociation constants (K_D_) were quantified by ITC. The data for the interaction of wild type Agno with the HP1α CSD shown in Fig. 2d and Extended data Fig. 1b and c are identical/from the same measurement.

**Extended data Fig. 2:**
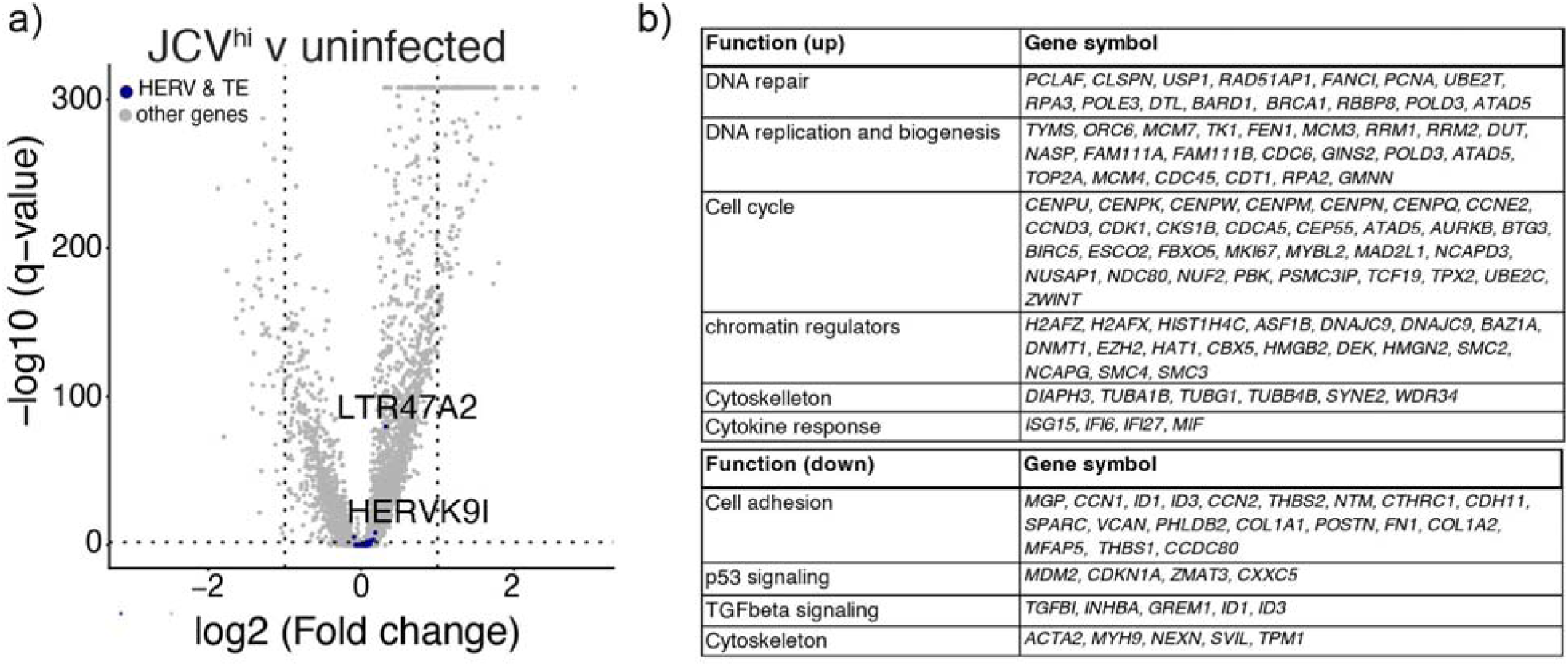
JCV infection does not derepress heterochromatin-silenced genes but induces a transcriptional program promoting DNA repair, replication and biogenesis and regulating cell cycle progression. a) JCV infection does not associate with de-repression of genes silence by heterochromatin such as hERVs and transposable elements (TE). Volcano plot showing the differential gene expression between JCV^hi^ cells and control cells. Genes encoding hERVs and TE that change in expression are indicated in blue. None of these genes changes more than 2-fold. b) Functional classification of the up- (top) or downregulated genes (bottom; 2-fold cutoff) in JCV^hi^ cells.

**Suppl. Table 1:**
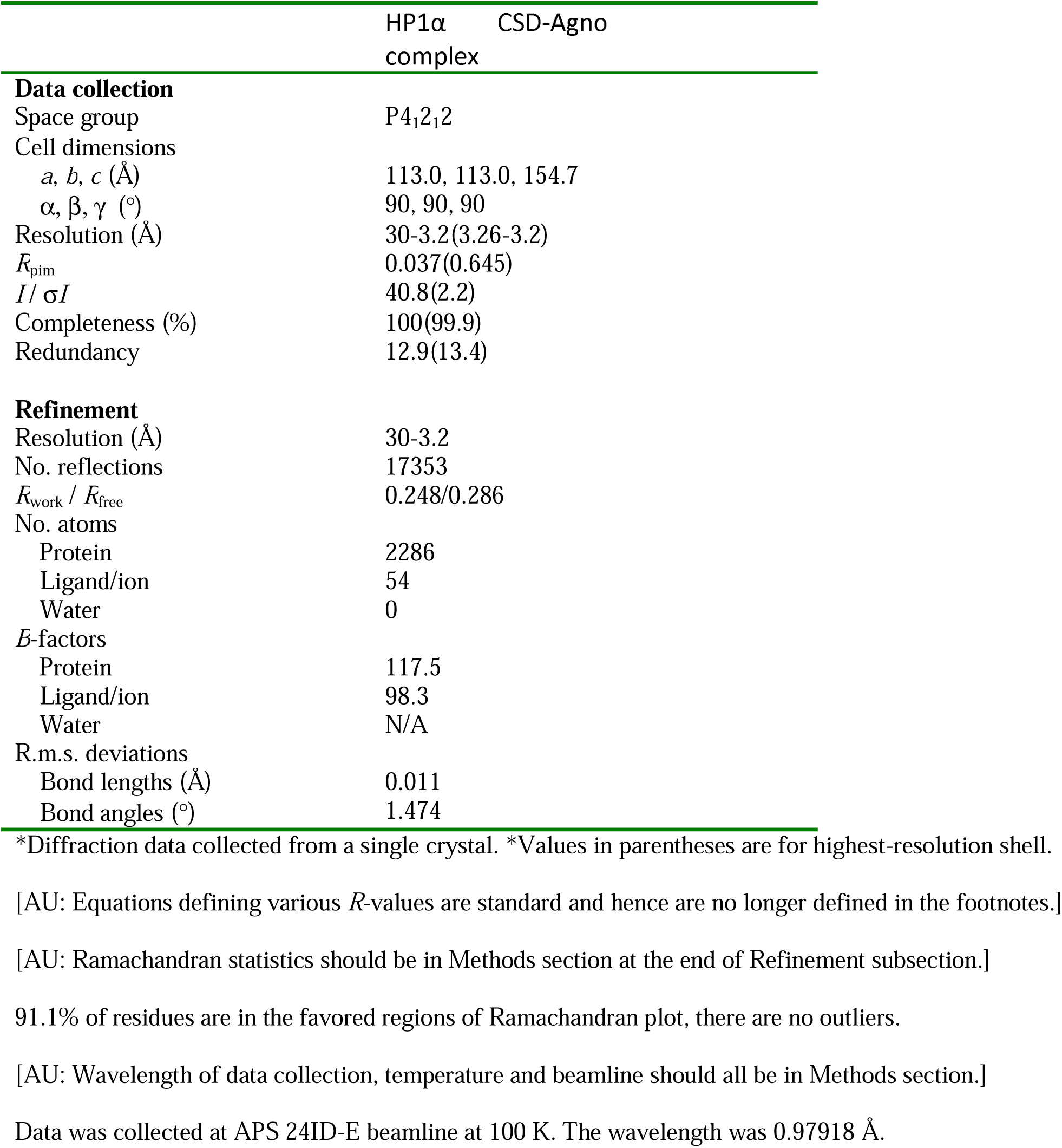
Data collection and refinement statistics (molecular replacement)

